# Viscoelastic properties of wheat gluten in a molecular dynamics study

**DOI:** 10.1101/2020.07.29.226928

**Authors:** Łukasz Mioduszewski, Marek Cieplak

## Abstract

Wheat *(Triticum spp*.) gluten consists mainly of intrinsincally disordered storage proteins (glutenins and gliadins) that can form megadalton-sized networks. These networks are responsible for the unique viscoelastic properties of wheat dough and affect the quality of bread. These properties have not yet been studied by molecular level simulations. Here, we use a newly developed *α*-C-based coarse-grained model to study ∼ 4000-residue systems. The corresponding time-dependent properties are studied through shear and axial deformations. We measure the response force to the deformation, the number of entanglements and cavities, the size of fluctuations, the number of the inter-chain bonds, etc. Glutenins are shown to influence the mechanics of gluten much more than gliadins. Our simulations are consistent with the existing ideas about gluten elasticity and emphasize the role of entanglements and hydrogen bonding. We also demonstrate that the storage proteins in maize and rice lead to weaker elasticity which points to the unique properties of wheat gluten.

## 1 Introduction

Wheat gluten is defined as the part of the wheat *(Triticum spp*.) flour that is insoluble in water. It can be obtained by gently washing away the soluble components (mostly starch and globular proteins) from the flour [1]. Over 75% of the insoluble mass consists of storage proteins [2]. They exhibit viscoelastic properties that are vital for the bread properties and quality [3]. Gluten proteins constitute less than 20% of the wheat dough mass, and yet it is those proteins that determine its elasticity [4]. Here, we neglect other substances present in real gluten samples and concentrate on the proteins.

Gluten proteins have sequences with repetitive low complexity fragments that are rich in glutamines and prolines [1]. As a result, they do not possess any established tertiary structure [2, 5, 6] and are thus (mostly) intrinsically disordered (IDPs). There are two kinds of gluten proteins: gliadins (around 70% of the total gluten protein mass) and glutenins [1]. The former are soluble in a 50% alcohol solution. Glutenins, on the other hand, may form covalent complexes through disulfide bonds [1] and are not affected by the alcohol.

The observed viscoelastic behavior of gluten implies the existence of properties that are both solid-like (an elastic response to deformation) and liquid-like such as the viscous drag that arises from irreversible deformations. These properties can be characterized by the dynamic Young modulus *G*\*. It is one of the few quantities that can be used to directly compare simulations to experiment. Here, we propose to use a one-bead-per-residue coarse-grained model [7, 8] that captures the most important features of gluten and agrees qualitatively with available data. The main limitation of the model is the usage of time scales that are substantially shorter than those used in the experiments. Therefore, we do not expect to match the actual experimental values, especially if they are frequency dependent. Our protein model enables milisecond-scale simulations of biochemically large protein systems, and preserves the amino acid specificity. Our molecular model allows us to determine the relative importance of different factors that affect the properties of gluten. This work is especially important because gluten proteins are insoluble and disordered [9] so one cannot infer the properties from any structural features. The only simulations of gluten proteins we could find considered single chains and not complexes [10, 11]. However, the crowding conditions that are present in gluten are expected to affect the single-chain behavior. We study systems of ∼ 4000 residues consisting of about 14 chains.

Existing theories of gluten have identified three main factors that contribute to its elasticity:

1. The formation of hydrogen and disulfide bonds: in every gluten protein over 30% of amino acids is glutamine, whose side chain can be both a donor and an acceptor of hydrogen bonds [3]. Disulfide bonds can hold gluten proteins together and generate a sort of polymer a gel [12];
2. The loops-and-trains model [2, 13] explains gluten elasticity by “loops” created by neighboring polymers: they stay together because of hydrogen bonding, but when some bonds are broken (e.g. by water), a free space emerges between the polymers. Under stretching, cavities disappear and hydrogen bonds are made between elongated chain fragments, forming “trains” that are parallel to the direction of stretching and cause resistance to further strain;
3. The entangled polymer network itself leads to strain resistance, regardless of its chemical nature [14].

We can study these phenomena in our molecular dynamics model by measuring the number of inter-residue contacts described by the Lennard-Jones (L-J) potential the number of disulfide bonds (that can form and rupture dynamically), the number and size of cavities (using the Spaceball algorithm [15, 16]) and the number of entanglements (using the Z1 algorithm [17-20]). Structural properties, like the distance between the ends of a protein chain, can be also evaluated. We have also computed the mechanical work required to stretch gluten.

Gluten proteins can be divided into two groups: long glutenins and short gliadins. The glutenins can easily form entanglements and inter-chain disulfide bonds, and are therefore thought to be the key ingredient of gluten elasticity [12]. The gliadins form fewer inter-chain disulfide bonds [1] and are thought to be largely responsible for the viscous properties of gluten [3].

We consider gliadins, glutenins and gluten (a mix of both) separately in order to elucidate the differences between these proteins. In addition, we also consider a control sample of proteins that do not exhibit such extraordinary viscoelastic properties [21]. Specifically, we use proteins derived from from maize and rice for this task. Thus we study 5 different systems. We show that many of the properties of gluten are substantially affected by “kneading” which we immitate by introducing periodic deformations.

## 2 Methods

For a detailed discussion of our model we recommend our other article [7]. Here, we will just describe it briefly. However, in our previous reports we did not discuss the dynamics of the disulfide-bond formation.

We use periodic boundary conditions in two directions (*X* and *Y*) and solid walls in the third direction *Z*. Our simulations cannot start from the native structure, because disordered proteins do not have one, so the starting conformations are constructed by generating self-avoiding random walks. They are then evolved by the methods of the coarse-grained molecular dynamics that were shown to correctly recreate properties of a set of intrinsically disordered proteins [7]. Rheological information is then obtained by deforming the box containing gluten proteins and recording the resulting force of response (see Subsection 2.2, “Simulation protocol”).

We simulate gluten as a set of protein chains in an implicit solvent. Each amino acid is represented by one pseudoatom and subsequent residues in a chain are connected by a harmonic potential. Chain stiffness is maintained through bond angle and dihedral angle potentials as obtained from a random coil library [7, 22]. These angle potentials have different forms based on the identity of residues constituting a given bond or dihedral angle: proline and glycine are treated as special cases. It is vital for gluten, which contains high amounts of those amino acids [1].

Electrostatics are governed by Debve-Hiickel potential with relative permittivity dependent on length [23]: *κ* = 40 nm^−1^ *r*. Other nonlocal interactions are modeled by 6-12 L-J potential, which can be attractive (in its full form) or repulsive (truncated at the minimum and shifted). Contacts between residues form and dissapear dynamically based on different criteria. The criteria have been developed by studying atomic-level overlaps that occur in contacts arising in structured proteins. We distinguish between contacts arising from sidechain-sidechain (ss), sidechain-backbone (bs) and backbone-backbone (bb) interactions. However, the resulting depth of the potentials well, *ϵ*, is common, no matter what is its origin. Contacts involving sidechains are amino-acid specific. Some gluten proteins are thought to be partially structured, so a part of the contacts are always attractive (see Subsection 2.1, “Coarse-grained model”).

### 2.1 Coarse-grained model

All details of the modeling are described in ref. [7] and here we focus only on some of them. We mark interacting pseudoatoms by indices *i* and *j*, 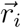 and 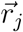 are their positions and 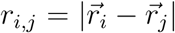 is the distance. Mass of each residue is set to the average amino acid mass *m*. The positions evolve according to the Langevin equations of motion with the time unit, *τ*, of ≈ 1 ns. The damping coefficient *γ* = 2*m****/****τ* (corresponding to overdamped dynamics [24]), and thermal white noise 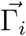 with variance *σ*^*2*^ = 2*γk*_*B*_*T:*

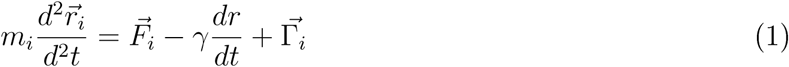

We solve the equations of motion by using the 5th order predictor-corrector algorithm [25]. Energy unit *ϵ* is of order 1.5 kcal/mol [26, 27]. The room temperature is then about 0.3 − 0.35 *ϵ/k*_*B*_ where *k*_*B*_ is the Boltzmann constant. Here, the simulations are run at 0.3 *ϵ/k*_*B*_. The results are presented, approximately, in the metric units.

The excluded volume of the residues (whether involved in a contact or not) is ensured by the L-J potential that is cut at *r*_*o*_ = 0.5 mm (so that *V*_*r*_ (*r*_*i,j*_ ≥ *r*_*o*_*) =* 0):

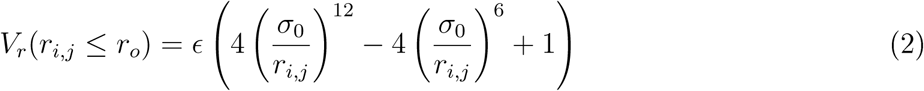

Where 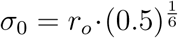, so that *V*_*r*_*(r*_*o*_) = 0. *i, j=i* + 2 interactions are always described by this repulsive potential. Connected *i, j*=*i* +1 beads interact via harmonic potential with the elastic constant of *k* = 5000 nm^-2^·*ϵ* and minimum at *r*_*b*_= 0.38 nm (consistent with other works [27, 28]).

The distinction between the bb, bs and ss contacts in our one-bead-per-residue model is accomplished in the following way. When two beads come sufficiently close to each other, we compute two vectors based on positions of three consecutive beads 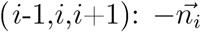 is the negative normal vector, approximately pointing in the sidechain direction, and the binormal vector *h*_*i*_ that points in the direction of a possible backbone hydrogen bond. The relative directions of these vectors are checked, and if, for instance, *h*_*i*_ and *h*_*j*_ point towards each other, a bb contact is made. Each type of amino acid has a specific number of contacts it can make with its virtual “backbone” and “sidechain”. Each ss contact making a pair of amino acids corresponds to different distances at the minimum of the L-J potential. On the other hand, that minimum is 0.5 nm for the bb contacts and 0.68 nm for the bs contacts. Independent of the source of the contact, the interaction is modeled by the potential of the same form:

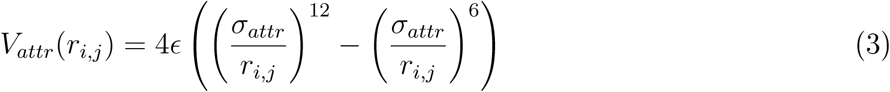

where *σ*_*attr*_ depends on the type of interaction. However, if the contact is due to the bb interactions, its strength is doubled. Doubling the depth of the L-J potential for bb contacts is consistent with the earlier literature findings [29]. If criteria for creation of a given type of contact are met, the attractive part of the potential is turned on at *r*_*min*_ in a quasi-adiabatically fashion (during 10 *τ*). When the beads are more than *r*_*f*_ = *f σ*_*attr*_ apart from each other it is turned off in the same way. This is why we call those contacts “dynamic” as they can appear and disappear during the simulation. In references [7, 30] and in this paper, we have used *f =* 1.5 (in an analogy to our description of relevant contacts in the structured proteins [26]). This model was previously used to study polyglutamine aggregation [30], so it should be well-suited for studying glutamine-rich gluten proteins. However, our more recent tests indicate that the choice *f* =1.3 is better. In particular, it is more consistent with the ANTON-based simulations of *α*-synuclein [31].

#### 2.1.1 Disulfide bonds

For modeling disulfide bonds, we considered using harmonic potential that is switched on and off, following Thirumalai et al [32]. However we have found out that it is equivalent but much simpler to use the L-J potential with the strength of 4 *ϵ* and 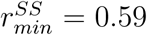 nm. We have checked that such a potential correctly recreates the experimental folding and unfolding times of crambin and ubiquitin (unpublished).

The disulfide bond potential is turned on and off in the same manner as other ss contacts, i.e. adiabatically within approximately 10 *τ*. It should be noted that the real timescales of the disulfide bond formation and their duration are much longer [33]. The only modification besides the amplitude change is that after two cysteins form such a contact, they cannot form additional disulfide bridges (without a prior rapture of the existing bridge).

#### 2.1.2 Ordered parts

Some of the gluten proteins contain domains near their termini that tend to be more structured than the whole chain. For example, homology of the A domain of high molecular weight glutenins to a plant inhibitor of digestive enzymes [34] suggests that one or two pairs of cysteines in that domain form fixed intra-chain disulfide bridges. We incorporated available information about possible structured parts of gluten by using a structure-based Go model for those regions [26]. The identification of segments that are recognized as structured domains was based on the UniProt [35] partition of gluten proteins into domains (see Table S1 for details). For the backbone stiffness of the ordered parts we used the same potential as for the unstructured parts.

The actual structures in the ordered parts were predicted by using the ITASSER. server [36, 37], then generated contact maps, based on those all-atom structures, using the overlap criterion (spheres made by heavy atoms of the residues in contact must overlap [26]). The contacts present in the contact map in the structured regions are always attractive (they are “static”, in contrast with the “dynamic” contacts operating in the disordered parts) and the L-J potential describing them has a minimum that corresponds to the distance between the *C*_*α*_ atoms in the reference all-atom structure.

Residues listed in the “static” contact map can still make “disordered” contacts with the residues outside the contact map. The number of “static” contacts made by a residue is determined by the same *r* <*r*_*f*_ criterion, but the potential does not switch off if *r* > *r*_*f*_.

### 2.2 Simulation protocol

We performed simulations of 5 different systems specified in Table 1 and described in more detail in Table S1. Storage proteins in the grains are separate [38]. They form large complexes only after the flour is mixed. This is reflected in our simulation protocol: we start simulations at a low density, give time for each chain to equilibrate separately [39] and then mix the chains together by shrinking the simulation box.

**Table 1:** The mean number of cysteines per chain *<n*_Cys_>, mean number of residues in a chain <*N*>, total number of residues Σ,*N* and the mean coordination number *z*, for all of the systems studied here. The last property was determined based on the simulations, the rest by the sequential make-up of the systems.

| system | $\langle n_{\text{Cys}} \rangle$ | $\langle N \rangle$ | $\Sigma N$ | $z$ |
| --- | --- | --- | --- | --- |
| maize | 6.1 | 221 | 3504 | $5.5 \pm 0.1$ |
| gliadins | 6 | 290 | 4322 | $6.88 \pm 0.03$ |
| glutenins | 7.2 | 445 | 3985 | $6.55 \pm 0.08$ |
| gluten | 6.5 | 384 | 4271 | $6.65 \pm 0.07$ |
| rice | 3.6 | 208 | 3503 | $5.62 \pm 0.09$ |

The first step of the simulations is generating the chains as self-avoiding-random-walks positioned randomly in a box with density *ρ* = 0.1 res/nm^3^ (residues per cubic nanometer). If an initial random walk configuration happens to cross the boundary of the box, the box is enlarged to keep all chains inside, resulting in a slighly lower initial density. The density of 0.1 res/nm^3^ is more than 30 times lower than the density of gluten, which should ensure that proteins are in monomeric or dimeric form during the initial equilibration (see inset 1 in Fig. 1). Boundary conditions along the *X* and *Y* axes are periodic, whereas the walls along the *Z* direction are repulsive. They are separated by the distance of *s*. The residues are repelled from the wall by the repulsive part of the L-J potential with depth 4*ϵ*.

**Figure 1:**
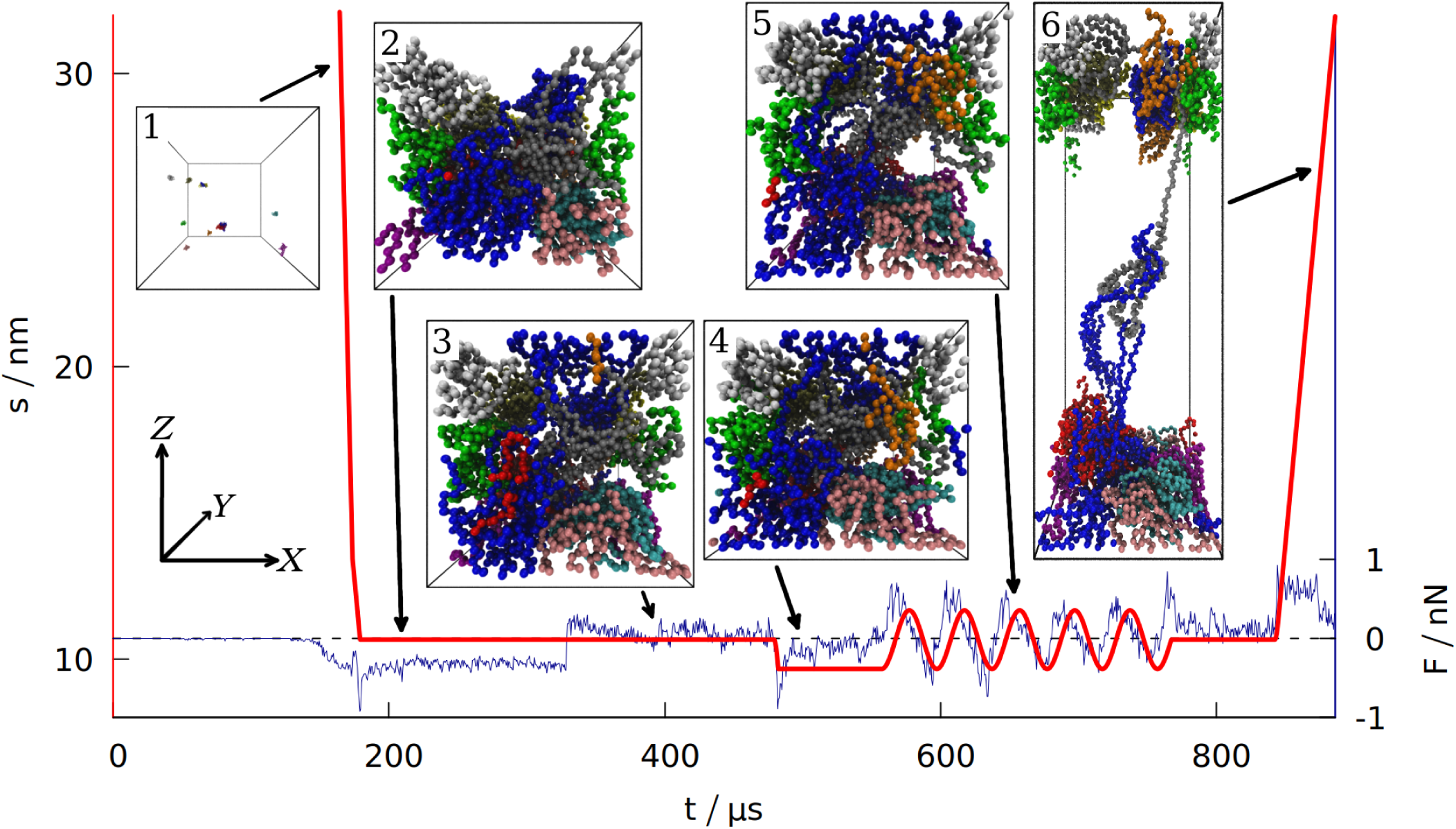
Time dependence of the distance *s* separating the two opposite box walls in the *Z* direction. The blue curve shows the force exerted on the system by those two walls (summed over all the residues and averaged over 100 ns time interval). The data is for gluten (defined in Table S1), with the density of 3.5 nm^-3^. The period of oscillations is 40 ms. The six insets correspond to the snapshots of the system taken at different stages of the simulation (each bead represents one amino acid, the protein chains are shown in different colors). The same coordinate system (top left) is used for all 6 insets.

After equilibrating the system for 125 *μ*s the box dimensions are reduced from the initial size with the speed of 2 mm/s. after the density *ρ*_0_ (usually corresponding to the real density of gluten [40] *ρ*_0_ ≈ 3.5 res/nm^3^) is reached (see Section 3 in the SI for more information about the density). At this stage, the distance between the repulsive walls has a value that is denoted by *s*_0_. The system is then equilibrated for another 150 *μ*s.

After the second equilibration stage (depicted in inset 2 in Fig. 1) the walls along the *Z* direction start to attract the proteins. When the distance between any residue and the wall gets closer than *d*_*min*_ = 0.5 nm, the attractive part of the L-J potential with depth of 4*ϵ* is turned on adiabatically (in the same manner as for disulfide bonds). Then the interaction between the wall and a residue attached to it is described by *V*_*wall*_ = 4*V*^*LJ*^*(r*_*i,w*_), where *r*_*i,w*_ is the distance between the *i*th pseudoatom and an *i*th interaction center on the wall. The *X* and *Y* coordinates of the *i*th interaction center are fixed at the values of the first bead attachment (came to the wall closer than *d*_*min*_ = 0.5 nm, which is the distance corresponding to the minimum of the *V*_*wall*_ potential). The *Z* coordinate of an interaction center is 0 on one rigid weall and *s* on the opposite wall. If a wall moves so that *s* ≠ *s*_0_, the interaction centers move together with the wall. If the bead departs away from the interaction center by more than 2 nm, it is considered detached and the interaction center disappears. The bead may reattach at another place later. For *d* < *d*_*min*_ the repulsive part of the L-J potential is always turned on to ensure that no residue can pass through the wall. The force acting on the walls is a sum of forces that individual residues exert on both walls. It is averaged over 100 ns interval to filter the thermal noise out. An alternative way to introduce interactions with the walls (involving the fcc lattice of psudoatoms) is discussed in Section 4 of the SI.

After the wall attraction is enabled, we give another 150 *μ*s for the equilibration of the system (inset 3 in Fig. 1). After this third equilibration stage, when the attraction is already turned on, the walls in the *Z* direction are moved to the distance of *s* = *s*_0_ − *A*, where *A* is the amplitude of the oscillations in the normal direction (we set *A* =1 nm). After being moved to the minimum (shown in inset 4 in Fig. 1), the wall positionchanges periodically: *s(t)* = *s*_0_ − *A cos (ωt)*. The wall at *Z*=0 stays fixed. In the experiments, *ω* ∼ 1 Hz [3], which is not possible to achieve in our simulations. Our default period is 40 *μ*s, corresponding to *f* ≈ 25 kHz. The maximum displacement is shown on inset 5 in Fig. 1.

For shearing simulations, the walls are also displaced with the same amplitude, *A*, but the oscillations take place in the *X* direction and are described by s*′* (*t*) = −*A* cos *(ωt)* (where *s*′ = 0 corresponds to one rigid wall being directly above the other). See Fig. S1 for an illustration.

After oscillations in the normal direction (or after additional 100 *μ*s of equilibration if no oscil-lations were made) the system is equilibrated again for 75 *μ*s and then the walls in the *Z* direction extend gradually to 2*s*_0_ so that total work required for such an extension can be measured, as well as the distance and force required to break the system apart (recreating conditions for measuring uniaxial extension [41]). Inset 6 in Fig. 1 depicts this elongation. The speed of elongation is 0.5 mm/s, which is a typical pulling speed employed in AFM stretching experiments in simulation [26, 42]. The maximum speed during oscillations (*v*_*max*_ = *Aω)* is even lower.

## 3 Results

### 3.1 Properties of gluten, maize and rice

Based on refs. [1, 4, 9, 12, 43-48] we chose proteins that were representative for each of the five systems we study: gliadins, glutenins, gluten and storage proteins from maize and rice. We then used the UniProt database [35] to select the corresponding sequences. The resulting compositions are described in Section 1 of the SI. Table 1 summarizes the key properties of each of the systems studied. A large number of cysteines per chain should increase the propensity to form inter-chain disulfide bonds. On the other hand, longer chains should make entanglements easier. Finally, the “stickiness” of the system is reflected in the coordination number ***z***, which is defined as the mean number of contacts made by each residue (both “dynamic” and “static” combined, see Subsection 2.1).

We performed three types of simulations: a) without any oscillations; b) with oscillations with deformation in the normal *(Z)* direction; c) with oscillations with deformation in the shear *(X)* direction. All of the properties are calculated in the stage after oscillations (in the absence of oscillations - after the equivalent time is passed). The parameter *z* does not seem to be affected by the oscillations.

However the values of 5 other quantities depend on the prior history of the system. Those quantities are: the number of entanglements *l*_*k*_ the maximum force F_max_, the mechanical work *W*_*max*_ required to stretch the system, the number of the interchain contacts *n*_*inter*_ and the RMSF (root mean square fluctuation) as averaged over all residues and determined as a deviation from a mean location of the residue.

It should be noted that each of the systems studied is simulated in a somewhat different box (to achieve the same density for different numbers of residues) and thus comparison of the absolute values of these quantities is not very meaningful. Instead, we analyze the ratios of the five quantities to their average values in the absence of oscillations (denoted by the tildas). Fig. 2 shows these ratios for the case of the shearing oscillations whereas Fig. S3 for the uniaxial oscillations.

**Figure 2:**
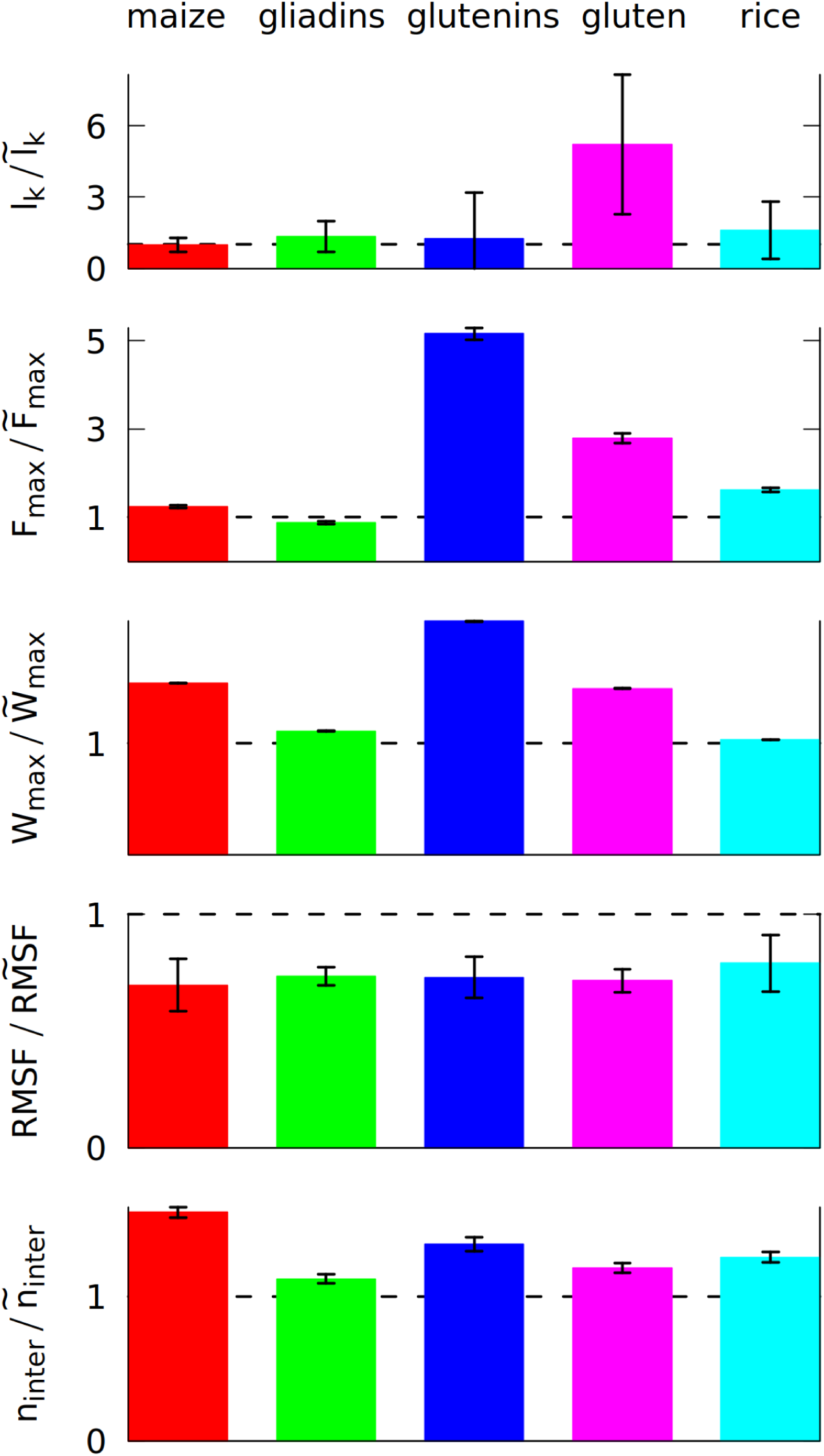
The ratio of 5 different properties calculated after 5 shearing oscillations (in the nominator), and without any oscillations (in the denominator, marked by ∼). The properties are the average number of entanglements *l*_*k*_, the maximum force F_max_ during the elongation in the last simulation stage and the maximum work W_max_ required to elongate the system in that stage, the number of inter-chain contacts n_inter_ and RMSF (root mean square fluctuation) averaged over all residues. The ratio of 1 is marked by a broken line. Each color corresponds to a different system, as displayed at the top, and defined in Table S1. The system density is 3.5 nm^-3^.

The first of the quantities studied is the total number of entanglements. We have determined *l*_*k*_ by using the Z1 algorithm [17]. An entanglement occurs if a path connecting the first and last residue in a chain cannot be shortened to a straight line without crossing a path from another chain [18, 19]. After minimizing the lengths of all such paths with the constraint that they cannot cross one another, each kink, where two paths intertwine, counts as an entanglement [20]. An example of an entanglement is shown in inset 6 in Fig. 1, where the end of the grey chain (a low molecular weight glutenin) crosses a loop made by a blue chain (a high molecular weight glutenin). Due to the low number of the chains, each snapshot of the simulation usually has only a few entanglements. We averaged *l*_*k*_ over all snapshots made after oscillations, and the simulations for each system were repeated at least three times with a different random seed (but the same set of seeds was used for the shearing, pulling and equilibrium simulations). The result shown in Fig. 2 uses the mean values from all simulations, but the error of the mean is quite high. Nevertheless, a clear incease is observed for gluten: the oscillating deformations cause more entanglements. We did not include self-entanglements (knots) in Fig. 2, but they are also present in gluten: some long proteins can form very deep knots (see Fig. S5), that can last for over 100 *μ*s.

The differences between the studied systems are most pronounced in the case of Fmax, the maximum force required to stretch the system in the last stage of the simulation (illustrated in inset 6 in Fig. 1). The blue curve in Fig. 1 shows how the force rises and then drops near the end due to the rapture. However, different random seeds may lead to different force curves for the same system. Fig. 3 shows three stretching force curves for three different simulations of gluten. Each curve has a number of force peaks. In order to determine the characteristic largest force, Fmax, we take 5 large force points and calculate their mean value. These mean F_max_ are also indicated in Fig. 3 as dotted lines surrounded by strips that represent their standard deviation. The F_max_ used in Fig. 2 is a weighted average of those values (from simulations with different random seeds), where the weights are calculated from the standard deviation of the mean. Large glutenins experience the largest, 5-fold increase in F_max_ after oscillations, while for gliadins there is even a very slight decrease. The rise in F_max_ for gluten is substantially bigger than for corn and rice proteins.

**Figure 3:**
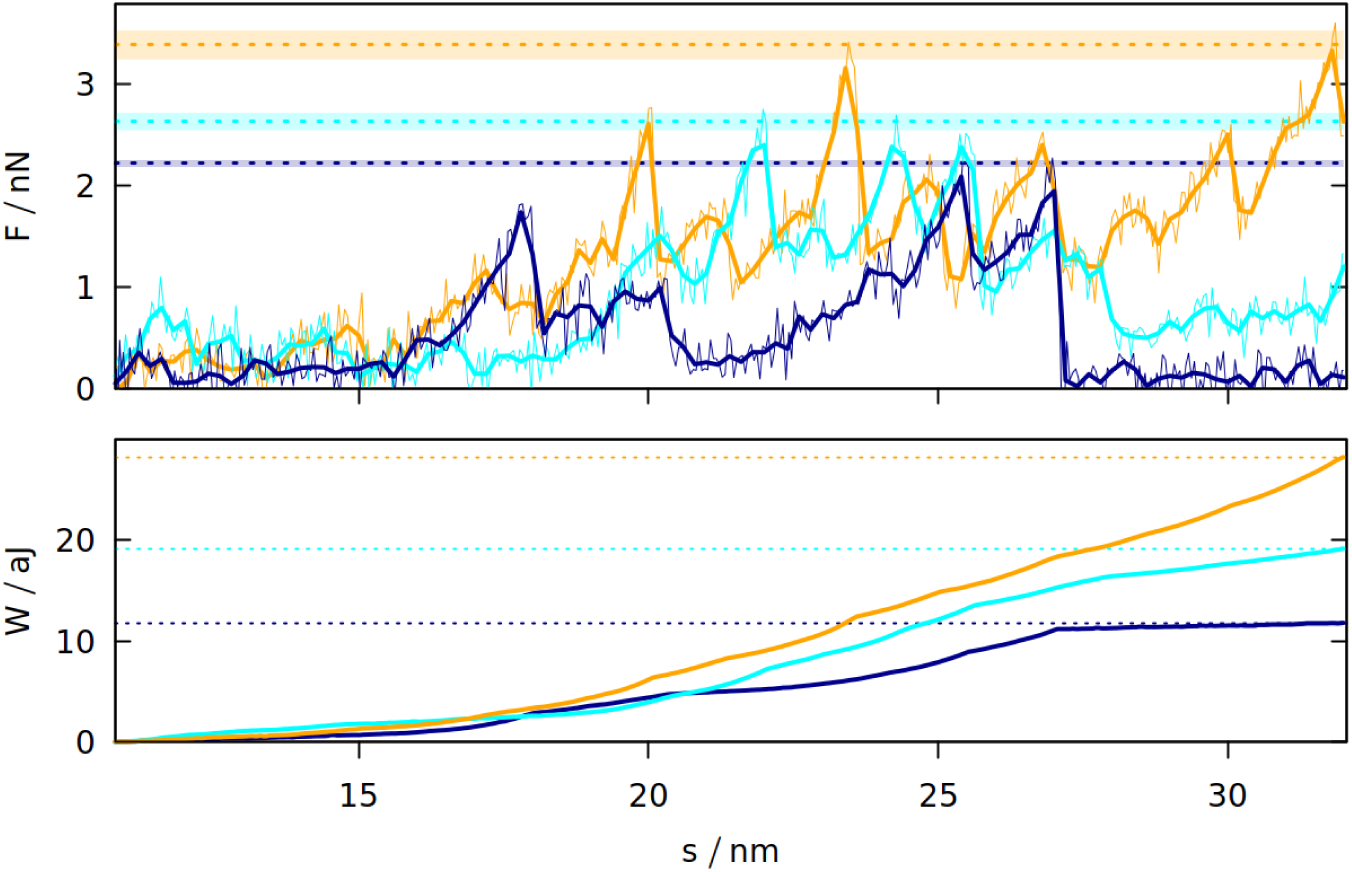
The force exerted on gluten with the density of 3.5 nm^-3^ (top panel) during elongation in the last stage of the simulation and the resulting work (bottom panel) as a function of the distance between the rigid walls. The three trajectories are shown in different colors. The dotted lines correspond to F_max_ arid W_max_ for a given trajectory. For F_max_ the strips around the dotted lines represent the uncertainty resulting from averaging F_max_ over 5 points with the biggest force. The oscillation period is 40 *μ*s. For *F*, the thin lines represent raw simulation data and the thick lines are smoothed.

The last stage of the simulations includes moving the Z walls apart with a constant speed. The response force may be easily integrated over distance, giving the work required to stretch the system. The maximum work W_max_ required to stretch the system is also bigger if oscillations were introduced before. Again, the difference is largest for glutenins, this time followed by maize proteins. Sometimes the force drops to near zero after reaching the maximum (meaning that the system was ruptured) and the work stops rising (as seen for the dark blue curves on Fig. 3). There are other cases where a substantial mechanical work is still required for elongating the system after reaching F max (cyan curve), but the total mechanical work W_max_ is correlated with the height of the F_max_ peak. The proteins apply some effective pressure on the walls, however it is much smaller than the mechanical stress resulting from the stretching (the atmospheric pressure would exert 0.01 nN of force on the walls).

Even if entanglements cause sharp peaks in the force curves, the resistance against deformation is also controlled by the number of inter-chain contacts, n_inter_, that have to be ruptured during elongation. The biggest increase in n_inter_ is observed for maize proteins. They have a lower number of contacts in total (as indicated by their coordination number ***z*** shown in Table 1), but their residue composition allows them to make more contacts due to the mixing effect of the oscillating deformations.

The oscillatory mixing effect is general and it operates in each of the systems studied. It results in a stronger network of proteins that are connected together more tightly, which results in the lowering of RMSF. Less mobile proteins may be more sturdy, but, as a trade-off, the maximum force may occur for a lower inter-wall distance s. We could not check (data not shown due to large errorbars) if this distance, averaged over simulations with different random seeds, depends on the system or on its prior periodic deformation. For the same reason, we do not show the increase in the number of inter-chain disulfide bonds.

### 3.2 The molecular explanation of gluten elasticity

According to the “loops and trains” picture, the response of gluten to deformation consists of two stages: at the first stage the chains elongate and the cavities between them shrink, which requires smaller force than disentangling the chains and moving them alongside each other that takes place in the second stage [13, 49]. We tested these predictions by measuring how 6 different quantities change during the final extension of the simulation box (see Fig. 4). The snapshots shown in this figure indicate that the “trains” do form, and that shearing between the chains drawn in red and blue may lead to a substantial force by disrupting and reforming the contacts between the chains. Indeed the number of inter-chain contacts decreases in the beginning of the elongation, but then it increases again as “trains” are formed. The chains become more rod-like and less globular, as indicated by the distortion parameter *w* [50] (averaged over all chains) that is equal to 0 for an ideal sphere, and 1 for an ideal rod. It is defined as 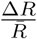, where 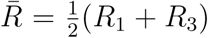 and 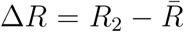; *R*_1_ is the smallest radius of inertia, *R*_*3*_ − the largest and *R*_*2*_ the intermediate.

**Figure 4:**
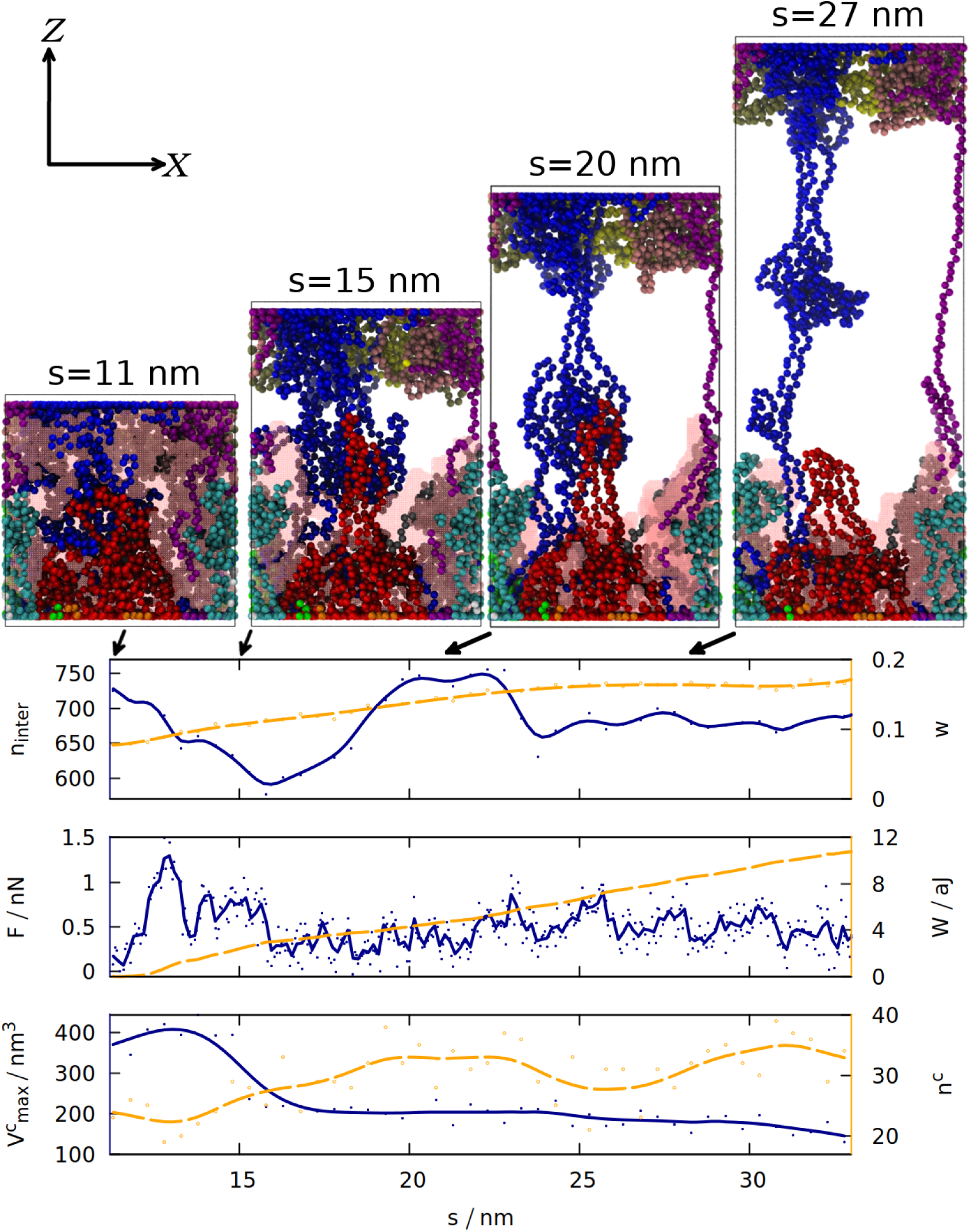
The time dependence of 6 different quantities that are measured for gluten (as defined in Table S1) during the elongation in the last stage of the simulation. The solid blue (orange) lines correspond to the quantities listed along the left (right) *y-axis*. The lines are smoothed, solid (hollow) dots represent raw data from simulation. The quantities analyzed are: the number of inter-chain contacts n_inter_ and the average distortion parameter *w*, force *F* and work *W*, the maximum volume of a cavity 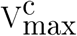 and the number of cavities n^c^. The 4 snapshots (similar to those on Fig. 1, but in the ortographic projection) correspond to the displacements *s* written above them and marked by the arrows. The same coordinate system (defined in the top left) is used for all snapshots. The biggest cavity is outlined by a pink mesh.

Formation of a “train” (two parallel chain fragments connected by hydrogen bonds) does not coincide with the maximum of the force, because F_max_ occurs near the beginning of the elongation and is associated with the biggest decrease in the number of contacts. Nevertheless, the force does not drop to zero and the work required to stretch the system steadily increases, because the chains from the opposite ends of the box are still connected and cause further resistance against deformation.

In order to quantify the number and size of cavities that form in gluten, we used the Spaceball algorithm [15, 16], which uses a probe with a user-defined radius to determine a three-dimensional contour of the system. The volume of the cavities is computed by filling a grid with those probes and rotating the grid with respect to the system. Our system is quite large, so we used grid with 0.1 nm lattice constant, a probe with radius 0.38 nm and one rotation of the grid. The snapshots generated show that the biggest detected cavity is dispersed over the whole simulation box, but during the elongation it becomes associated with the set of proteins attached to the *Z* = 0 wall (for the trajectory shown). As they are more and more elongated, the size of the cavity decreases. Larger cavities become divided into many smaller cavities, which is reflected in the increase in *n*_*c*_.

Fig. 4 illustrates just one simulation (the same properties for the same system, but with a different random seed, are shown in Fig. S6), but the volume of the biggest cavity indeed decreases after elongation in most cases (see Fig. S4). The chains become more rod-like not only during the final stretching, but also after the oscillations (see Fig. S2): deformations cause the proteins to be briefly stretched, but after the deformation ends, the proteins do not return to the exactly the same shape as before, but stay elongated in one direction (which may be called a memory effect).

### 3.3 Determining the dynamic shear modulus

One of the few fundamental (independent of the size and shape of the sample) rheological properties of gluten is the dynamic shear modulus *G* = G*^*′*^ + *iG*^″^, measured by deforming the gluten sample with sinusoidal deformation (shear strain for *G\** and normal strain for *E\**) [41]. The real part (*G*^*′*^, elastic modulus) represents the in-phase part of stress response, the imaginary part (*G*^″^, loss modulus) the out-of-phase response (phase difference *δ* is defined as tan 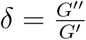). This is a non-destructive method, because the amplitude of oscillations is small (usually up to a few percent [3]). The total force *F*(*t*) that the system under a shear deformation exerts on the walls in the *X* direction (see Fig. S1), divided by the surface area of a wall *S*, defines shear stress *ϕ(t) = F*_*X*_*(t)/S* (the usual symbol for shear stress, *τ*, is a time unit in our model, so we use *ϕ* instead). A strain defined as *γ* = *γ*_0_ cos (*ωt*) is expected to induce a stress in a similar form, shifted in phase by *δ: ϕ* (*t*) = *ϕ*_0_ cos (*ωt* + *δ)*. Dynamic shear modulus *G\** = *G*^′^ + *iG*^″^ is then defined as [51]:

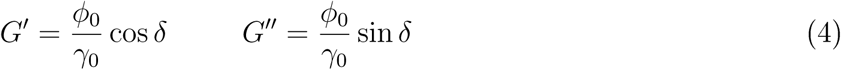

where *ϕ*_0_ and *γ*_0_ are the amplitudes of stress and strain accordingly. We define it by fitting to functions *s*^′^ *(t)* = *A* cos *(ωt)* and *F(t) = F*_0_ cos *(ωt* + *δ*). Here *δ* is the phase difference, *γ*_0_ = *A/s*_0_ and *ϕ*_0_ = *F*_0_/*S*. This is a very basic method of determining the shear modulus and more advanced methods can be used [52].

In Table 2 and Fig. 5 we compare the results with experiment. Table 2 shows that gliadins are responsible for the viscous properties of gluten (big *δ*), and glutenins for the elastic part (small *δ*). Despite largely different values of the results, this trend is still visible in our simulations for the longest oscillation period (70 *μ*s): tan*δ*_sim_ is biggest for gliadins and smallest for glutenins.

**Table 2:**
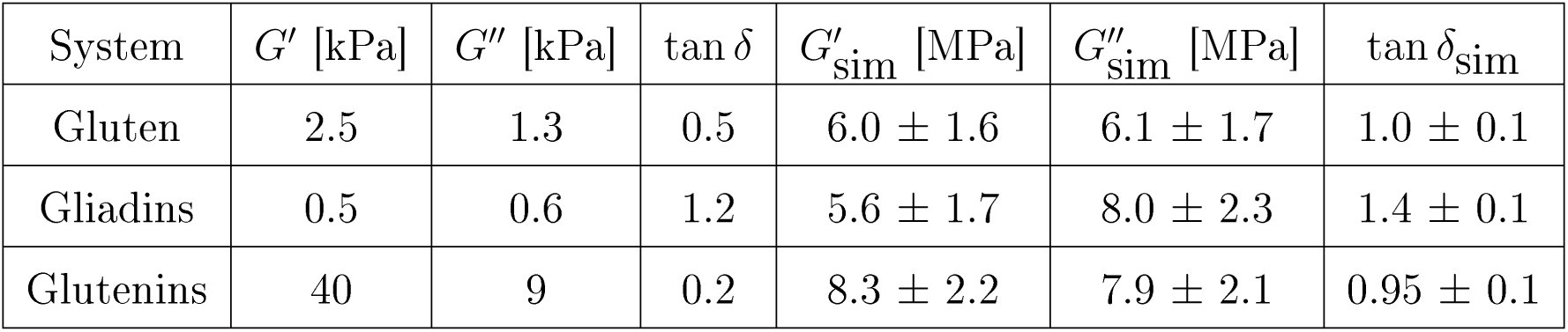
Shear modulus of dried gluten and its fractions, found by experiment with oscillation frequency *f* = 1 Hz [3], and by our simulations (marked by subscript _sim_) with *f ≈* 14 kHz (corresponding to the longest oscillation period we used, 70 *μs*).

**Figure 5:**
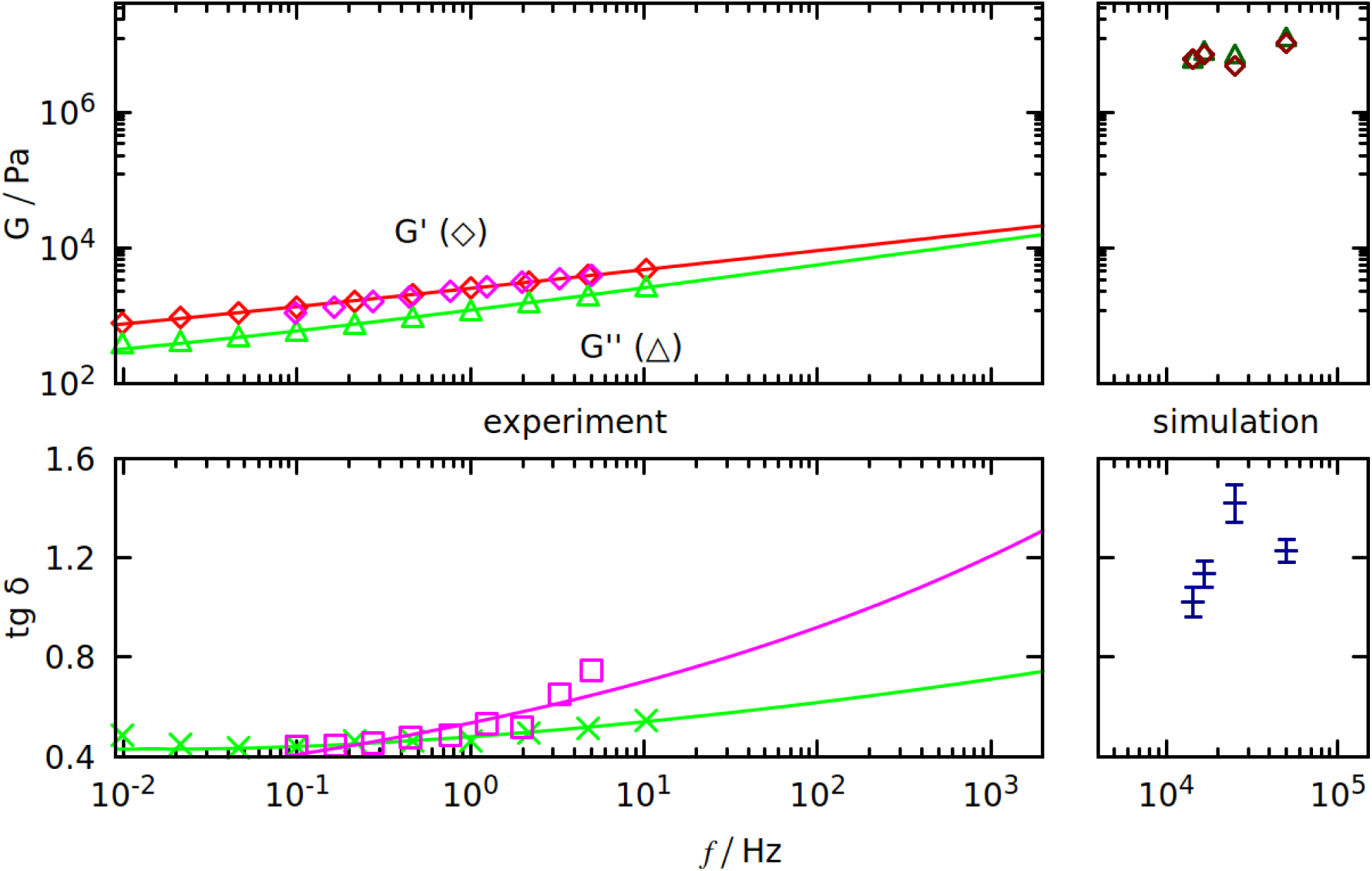
Dynamic shear modulus (top panel) and tan *δ* (bottom panel) as a function of oscillation frequency *f*. Results are from our simulations (on the right) and from experiments (on the left, purple points are from [54], red and green from [53]). All curves fitted to the data are of the form *a* · *f*^*b*^ + *c*, where *a, b* and *c* are the fit parameters.

We use frequencies that are 4 orders of magnitude larger than in experiment, but some extrapolations can be deduced from the experimental frequency spectra of gluten’s *G\**. We used the results from two experimental papers [53, 54]. Both predict that tan *δ* should be rising with frequency - see the bottom panel of Fig. 5, where tan*δ*_sim_ is between the extrapolations predicted by [53] (green curve) and [54] (purple curve). Such large discrepancies are the effects of different gluten compositions, that greatly influence the results (we use the Aubaine dataset from [54]).

Growing *δ* means that *G*^*′*^ and *G*^″^ converge, as seen on the top panel of Fig. 5. Both *G*^′^ and *G*^″^ are rising with *f*, however this rise is insufficient to explain why the values from simulation are 3 orders of magnitude larger. Either there is a significant non-linearity for higher frequencies and the extrapolation no longer holds, or this is a limitation of our model. We use implicit solvent which cannot account for differences in gluten’s hydration that may greatly affect the results (even,, dry” gluten is slightly hydrated) [53].

On the other hand, the results of the simulation are in good agreement with the experiments done on gluten-based bioplastics, where gluten proteins are mixed with glycerol and melted at over 380 K [55]. Even for bioplastics, the tensile strength (the stress needed to break the deformed system) is less than 1 MPa [55], while the F_max_ divided by the wall area (which is around 100 nm^2^) can be more than 10 times higher (see Fig. 3).

## 4 Discussion and conclusions

We have used our coarse-grained model with the implicit solvent to provide microscopic-level understanding of gluen and the related proteins. Kneading and mixing gluten samples is known to increase their tensile strength [56]. In our simulation, we mimick these actions by introducing periodic deformations. We have found that as a result of these deformations gluten and glutenins became significantly more resistant to stretching, as indicated by the rise in F_max_ and W_max_ Our model is also able to correctly distinguish gluten from storage proteins from other plants, and to tell the difference between durable glutenins and viscous gliadins. This difference appears to come from the combined effect of entanglements and inter-chain contacts (that represent mainly hydrogen bonds between glutamines in the case of gluten). Maize proteins formed more inter-chain contacts than rice proteins (which may justify why maize flour is more useful for making gluten-free doughs [57]). Its chains were too short to gain additional mechanical resistance from being entangled.

The importance of entanglements and inter-chain hydrogen bonding is already well-known in the experimental literature [12, 14], but our simulations provide an additional confirmation of that. We also elucidated microscopic mechanisms of the loops-and-trains picture [13]. We have found that the decrease in the volume of the biggest cavity is accompanied by the rise in the number of the inter-chain contacts during stretching.

Due to the small system size only a few inter-chain disulfide bonds were simultaneously present during our simulations (most of them were observed for maize proteins) and they did not seem to influence the results significantly. It is possible that they may become more important at larger scales, where covalent linkage of the chains affects properties like solubility [1]. However, for about 20 gluten chains, the entanglements and non-covalent contacts are sufficient to explain elasticity of the system.

We used the periodic deformation to calculate the dynamic Young modulus *G\**. We have observed that the theoretical frequency dependence shown in Fig. 5 agrees with the experimental results for glycerol-plasticized gluten [55] better than with those for unrefined hydrated gluten [53, 54]. In our implicit solvent model glycerol may theoretically replace water as the solvent. However, our model was parameterized with the assumption of a watery solvent [7]. It should be noted, however, that the experimental range of the frequencies is orders of magnitude smaller than implemented here. Further simplifications are needed to access the longer time scales.

Another reason for the very high *G\** may be the coarse-graining itself: a coarse-grained pseudoatom has smaller volume than an all-atom amino acid. This may lead to diffrent elastic behavior, as was observed previously in the case of virus elasticity [58].

Despite the deciding role of gluten in determining the dough quality, its measured *G\** values are not strongly correlated with it [3]. During baking, the gas bubbles inside the dough extend it far beyond the limit of linear deformations. Therefore measurements of extension to over 200% of the initial size are more informative [3]. Such extensions were generated in the last stage of our simulations and were discussed above. Unfortunately, properties like W_max_ and F_max_ depend on the system size and geometry, so they cannot be directly compared to experimental values.

Another hypothesis that can be checked in our simulation is that gluten proteins can form a *β*-spiral from consecutive *β*-turns made by repetitive fragments like QPGQ [11] (such fragments are common especially in glutenins [2]). Scanning tunneling microscopy discovered a groove with 1.5 nm periodicity in glutenins [2]. However, the sample preparation procedures used prevented the study from validating the existence of the *β*-spiral in bulk gluten. Models of single glutenin structures consisting of repetitive *β* [59] and *γ* [10] turns were reported in the literature, however we did not observe similar structures in our simulations.

We do not discuss here how gluten proteins interact with the human body, but this widely disputed subject [60, 61] can also be tackled by our model in the future. Simulations can also check what part of gluten proteins is solvent-accessible, and thus available for digestive enzymes in the stomach, which could shed new light on the gluten intolerance problem [62].

## 5 Acknowledgements

This research has been supported by the National Science Centre (NCN), Poland, under grant No. 2014/15/B/ST3/01905 and the European H2020 FETOPEN-RIA-2019-01 grant PathoGelTrap No. 899616. The computer resources were supported by the PL-GRID infrastructure. We thank Mariusz Raczkowski for his help in developing the potential for the disulfide bonds.

## Notes

### Competing Interest Statement

The authors have declared no competing interest.

