## Supplementary Information for "Viscoelastic properties of wheat gluten in a molecular dynamics study"

### Viscoelastic properties of wheat gluten in a molecular dynamics study - Supplementary Information

#### Contents

**1 Composition of gluten and its fractions** **1**

**2 Supplementary figures** **4**

**3 Choosing the density** **7**

**4 Choosing the wall interaction model** **8**

#### 1 Composition of gluten and its fractions

We tried to accurately recreate proportions of different gliadin and glutenin types in gluten (after [1] and [2]). Gluten composition depends on the wheat cultivar [3]. We chose the cultivars with the biggest amount of glutenins, because they are more important for gluten elasticity [4]. All sequences were downloaded from the UniProt database [5]. We excluded from them the signaling sequences. The number of residues in a single chain is a sum of the unstructured and structured parts (in bold). Sum with two bold terms indicates two structured domains.

In addition to gluten, gliadins and glutenins we also investigated properties of storage proteins from maize (*Zea mays*) [6–8] and rice (*Oryza sativa*) [9–11].

It is important to note that one gliadin $\gamma$ that we tried to use (UniProt code P04729) produced significantly different results from other gliadins and differed in sequence. We do not know what caused the discrepancies, so we decided not to use P04729 in our simulations.

*

**Gluten**

| Protein type (gluten) | Uniprot code | Number of chains | Number of residues $N$ | Total number of res. $\Sigma N$ |
|---|---|---|---|---|
| LMWGS, type 1D1 | P10386 | 3 | 114+**170** | 852 |
| HMWGS, type DX5 | P10388 | 1 | **89**+696+**42** | 827 |
| HMWGS, type DY10 | P10387 | 1 | **119**+466+**42** | 627 |
| Gliadin $\alpha/\beta$, type MM1 | P18573 | 3 | 130+**157** | 861 |
| Gliadin $\gamma$, type B | P06659 | 3 | 117+**155** | 843 |
| Gliadin $\omega$, type 1,2 | Q9FUW7 | 1 | 261 | 261 |
| Sum | | 12 | | 4271 |

**Glutenins**

| Protein type (glutenins) | Uniprot code | Number of chains | Number of residues $N$ | Total number of res. $\Sigma N$ |
|---|---|---|---|---|
| LMWGS, type 1D1 | P10386 | 6 | 114+**170** | 1704 |
| HMWGS, type DX5 | P10388 | 2 | **89**+696+**42** | 1654 |
| HMWGS, type DY10 | P10387 | 1 | **119**+466+**42** | 627 |
| Sum | | 9 | | 3985 |

**Gliadins**

| Protein type (gliadins) | Uniprot code | Number of chains | Number of residues $N$ | Total number of res. $\Sigma N$ |
|---|---|---|---|---|
| Gliadin $\alpha/\beta$, type MM1 | P18573 | 7 | 130+**157** | 2009 |
| Gliadin $\gamma$, type B | P06659 | 6 | 117+**155** | 1632 |
| Gliadin $\omega$, type 5 | Q402I5 | 1 | 420 | 420 |
| Gliadin $\omega$, type 1,2 | Q9FUW7 | 1 | 261 | 261 |
| Sum | | 15 | | 4322 |

Table S1, part 1: composition of gluten, glutenin and gliadin systems used during the simulations. Total number of residues ($\Sigma N$) is the number of residues in a single chain ($N$) multiplied by the number of chains. If structured domains are present, the number of residues in a single chain is a sum of the unstructured and structured parts (in bold).

**Maize**

| Protein type (maize) | Uniprot code | Number of chains | Number of residues $N$ | Total number of res. $\Sigma N$ |
|---|---|---|---|---|
| $\alpha$-zein, type 4 | O48966 | 3 | 245 | 735 |
| $\alpha$-zein, type 16 | P04700 | 4 | 242 | 968 |
| $\alpha$-zein, type 19C2 | P06677 | 3 | 219 | 657 |
| $\gamma$-zein | P08031 | 2 | 164 | 328 |
| Glutelin-2 | P04706 | 4 | 204 | 816 |
| Sum | | 16 | | 3504 |

**Rice**

| Protein type (rice) | Uniprot code | Number of chains | Number of residues $N$ | Total number of res. $\Sigma N$ |
|---|---|---|---|---|
| Glutelin, type-A 1 | P07728 chain 1 | 2 | 282 | 564 |
| Glutelin, type-A 1 | P07728 chain 2 | 2 | 193 | 386 |
| Glutelin, type-A 3 | Q09151 chain 1 | 1 | 281 | 281 |
| Glutelin, type-A 3 | Q09151 chain 2 | 1 | 191 | 191 |
| Glutelin, type-B 1 | P14323 chain 1 | 2 | 278 | 556 |
| Glutelin, type-B 1 | P14323 chain 2 | 2 | 197 | 394 |
| Glutelin, type-D 1 | Q6K508 chain 1 | 1 | 258 | 258 |
| Glutelin, type-D 1 | Q6K508 chain 2 | 1 | 199 | 199 |
| Prolamin 14E | Q0DJ45 | 1 | 131 | 131 |
| prolamin C | P17048 | 1 | 137 | 137 |
| 10 kDa prolamin | Q0DN94 | 1 | 110 | 110 |
| Prolamin 14P | Q42465 | 1 | 132 | 132 |
| 19 kDa globulin | P29835 | 1 | 164 | 164 |
| Sum | | 17 | | 3503 |

Table S1, part 2: composition of rice and maize systems used during the simulations. Total number of residues ($\Sigma N$) is the number of residues in a single chain ($N$) multiplied by the number of chains.

#### 2 Supplementary figures

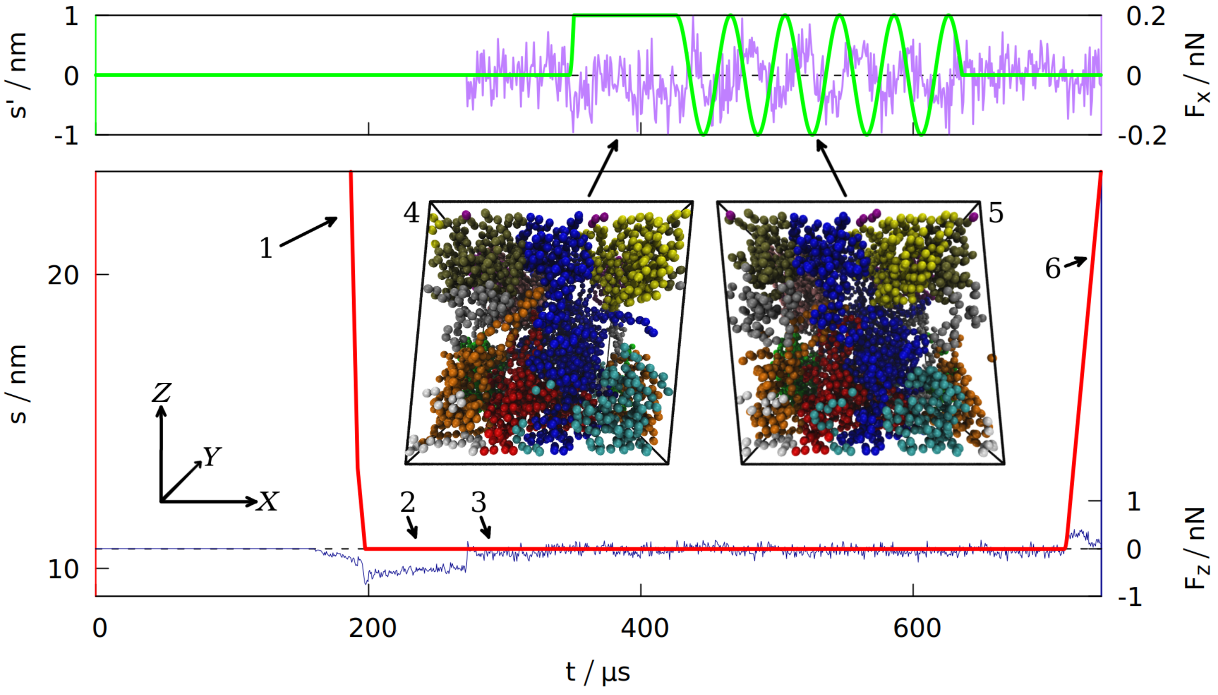


Figure S1: Time dependence of the wall displacement $s'$ in the $X$ direction (top panel) and the distance $s$ separating the two opposite box walls in the $Z$ direction (bottom panel). The purple curve (top panel) shows the $X$ component of the force exerted on the proteins by the walls (summed over all the residues and averaged over 100 ns time interval), the blue curve (bottom panel) shows the $Z$ component of that force. The data is for gluten (defined in Table S1), with the density of 3.5 $nm^{-3}$. The period of oscillations is 40 ms. The numbers with arrows correspond to the stages of the simulation shown on Fig. 1 in the main article. Snapshots 4 and 5 show the system during shearing oscillations (each bead represents one amino acid, the protein chains are shown in different colors). The coordinate system (top left) is the same as in Fig. 1.

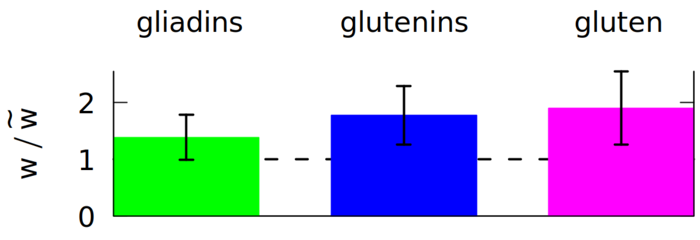


Figure S2: The ratio of the average distortion parameter $w$ [12] calculated after 5 shearing oscillations (in the nominator), and without any oscillations (in the denominator, marked by ˜). Period of oscillations is 70 $\mu$s (lower periods did not result in such significant changes in $w$). The ratio of 1 is marked by a broken line. Each color corresponds to a different system, as displayed at the top, and defined in Table S1. The system density is 3.5 $nm^{-3}$.

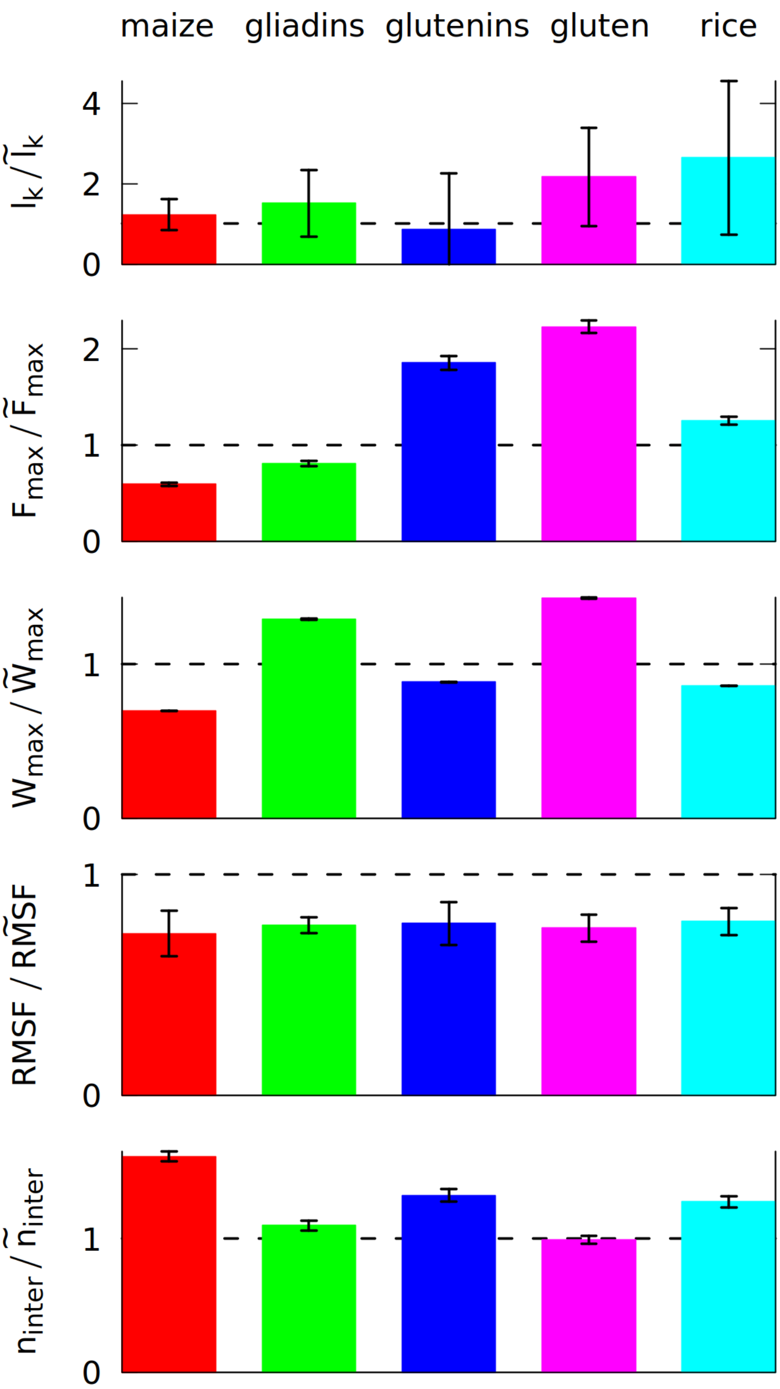


Figure S3: The ratio of 5 different properties calculated after 5 axial oscillations (in the nominator), and without any oscillations (in the denominator, marked by ˜). The properties are the average number of entanglements $l_k$, the maximum force $F_{max}$ during the elongation in the last simulation stage and the maximum work $W_{max}$ required to elongate the system in that stage, the number of inter-chain contacts $n_{inter}$ and RMSF (root mean square fluctuation) averaged over all residues. The ratio of 1 is marked by a broken line. Each color corresponds to a different system, as displayed at the top, and defined in Table S1. The system density is 3.5 $nm^{-3}$.

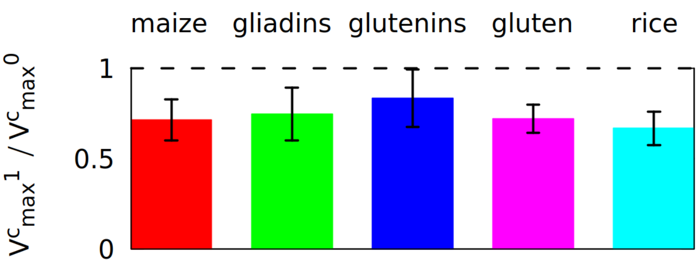


Figure S4: A ratio showing a decrease in the volume of the biggest cavity during the last stage of the simulations (elongation). The $V^c_{max}{}^0$ is the average taken over the first half of the elongation process, $V^c_{max}{}^1$ is the average taken over the last half. The ratio of 1 is marked by a broken line. Each color corresponds to a different system, as displayed at the top, and defined in Table S1. The system density is 3.5 nm$^{-3}$.

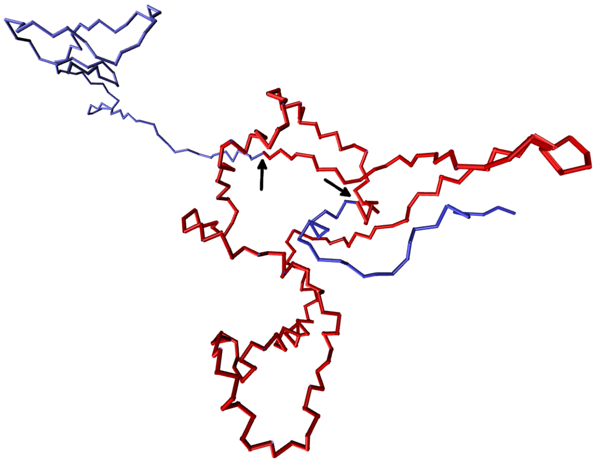

Figure S5: An example of a knotted high molecular weight (HMW) glutenin (from a gluten simulation with $\rho = 4$ nm$^{-3}$). Knotted area is shown in red, knot ends are indicated by black arrows.

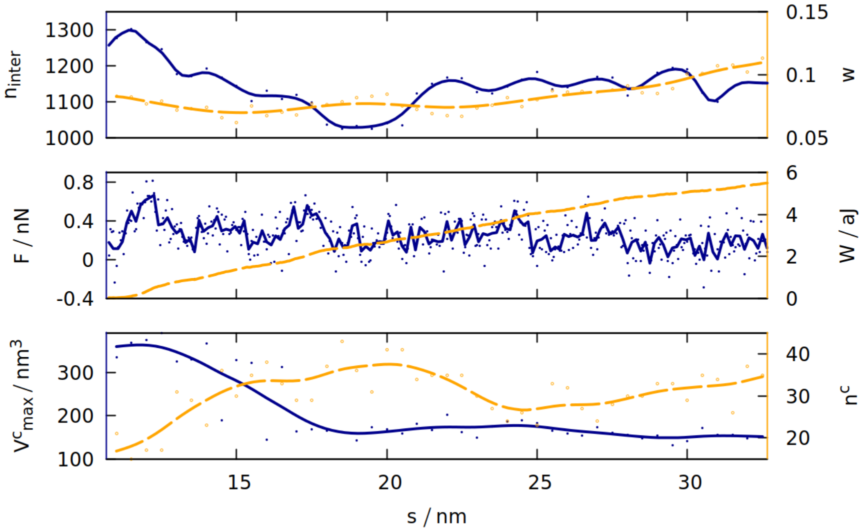


Figure S6: The time dependence of 6 different quantities (defined for Fig. 4 in the main article) that are measured for gluten (defined in Table S1) during the elongation in the last stage of the simulation. The solid blue (orange) lines correspond to the quantities listed along the left (right) $y$-axis. Solid (hollow) dots represent raw data from simulation.

### 3 Choosing the density

Real samples of gluten usually contain water, so in our implicit solvent model the gluten density may be higher than it should be. However in our one-bead-per-residue model we do not have an explicit representation of the sidechains, so each residue occupies less space than in reality. Therefore we have checked that the density $\rho_0 \approx 3.5$ res/nm$^3$ is indeed correct by comparing density profiles for various $\rho$. $\rho_0$ is near the threshold where the density profile becomes homogenous (see Fig. S7).

We studied densities ranging from 2 nm$^{-3}$ to 4 nm$^{-3}$, to find if the experimental $\rho_0$ =3.5 $nm^{-3}$ density is indeed correct for a system with implicit solvent. As the density rises, there is less free space, so the volume of the biggest cavity should decrease. However Fig. S8 shows that this volume rises at first: this is due to the fact that the density 2 $nm^{-3}$ is too low and there are free spaces in the simulation box not occupied by proteins (Spaceball distinguishes free space from a cavity inside a protein system). $V^{c}_{max}$ starts to decrease only for densities $\rho > \rho$ =3.5 $nm^{-3}$, so $\rho_0$ =3.5 $nm^{-3}$ is indeed the correct density (the system is homogenous, but there is still space for cavities).

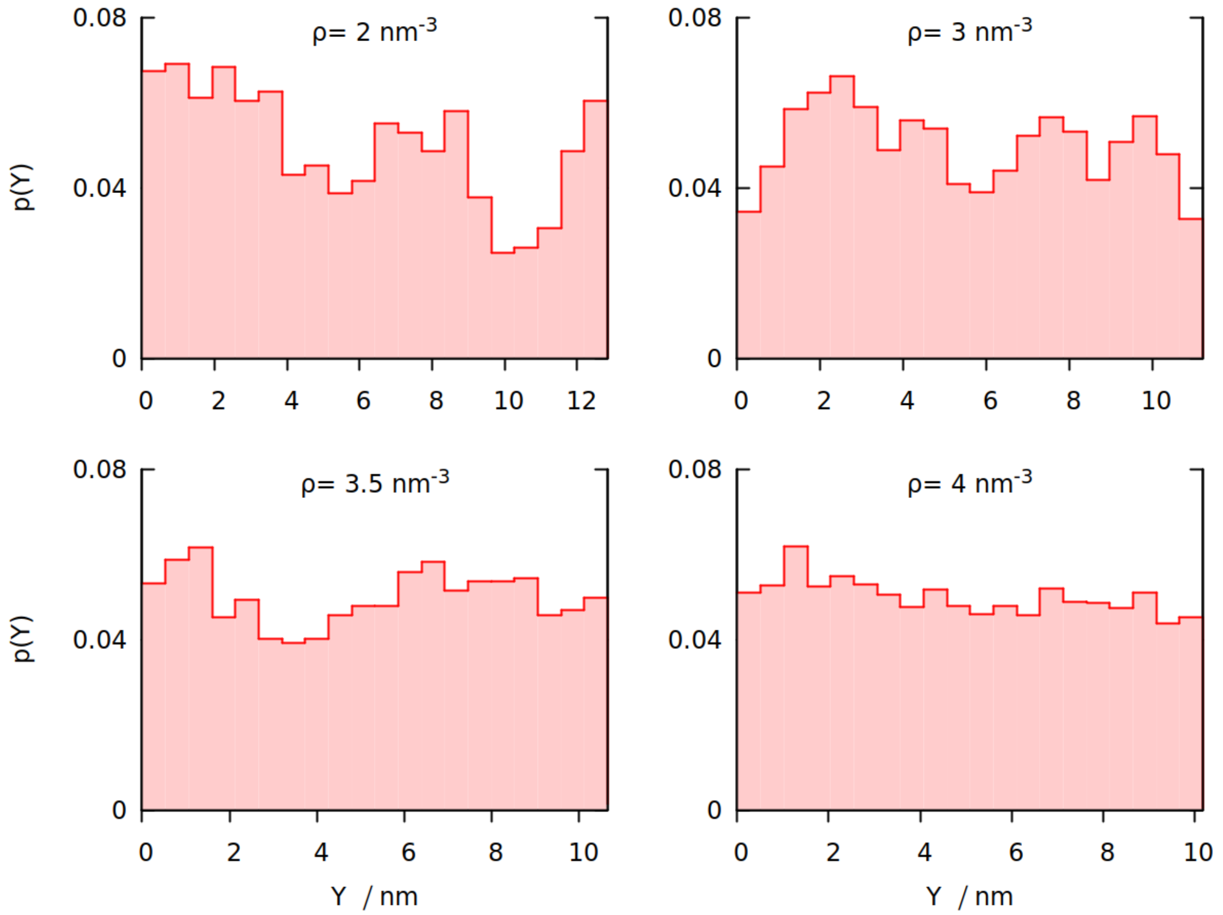


Figure S7: Density profiles along the $Y$ axis, summed over $X$ and $Z$. The profiles are snapshots from a simulation of gluten (with densities specified on the picture), taken in the stage after periodic uniaxial oscillations with period 20 $\mu$s.

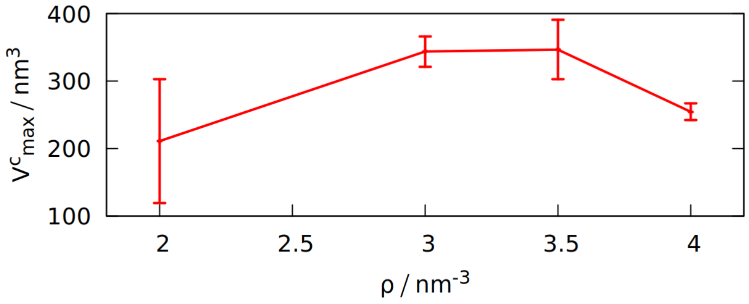


Figure S8: Density dependence of the maximum volume of a cavity, $V^c_{max}$, computed after oscillations with period 20 $\mu$s for gluten.

### 4 Choosing the wall interaction model

The way of attaching residues to the walls is crucial for gathering information about elastic properties of the system. Attaching them harmonically caused numerical instabilities and made it impossible for a residue to leave the wall. Therefore we compared two different methods: a) interaction centers put on an fcc lattice with lattice parameter 3 Å; b) dynamic interaction centers that appeared adiabatically when a residue approached the wall. In both methods the centers interacted via Lenard-Jones potential with strength 4 $\epsilon$, the same as for disulfide bonds. The FCC wall consisted of two layers of beads cut through the [111] crystallographic direction.

FCC method requires more computational power in order to simulate all the wall beads, and it results in stronger attraction: in the adaptive method a protein residue is always attracted by just one center, in FCC walls nearby residues also attract the residue. Density profiles along the Z axis (Fig. S9) show that for the FCC walls residues attracted to the walls gather close to them, making the profile non-uniform. This is not observed for the „adaptive" walls.

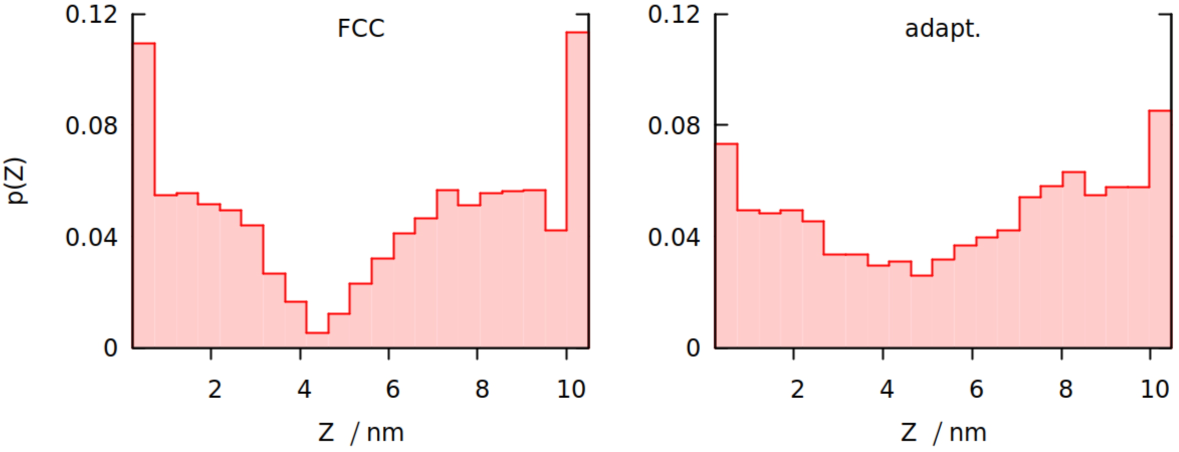


Figure S9: Density profile along the $Z$ axis, summed over $X$ and $Y$. The profiles are snapshots from a simulation of gluten (with densities specified on the picture), taken in the stage after periodic uniaxial oscillations with period 20 $\mu$s.

The force response for shearing is much closer to a sinusoid shape for adaptive walls (see Fig. S10), because FCC walls can cause „slipping" of residues from one interaction center to another. For the above reasons we decided to use the adaptive walls for determining gluten elasticity.

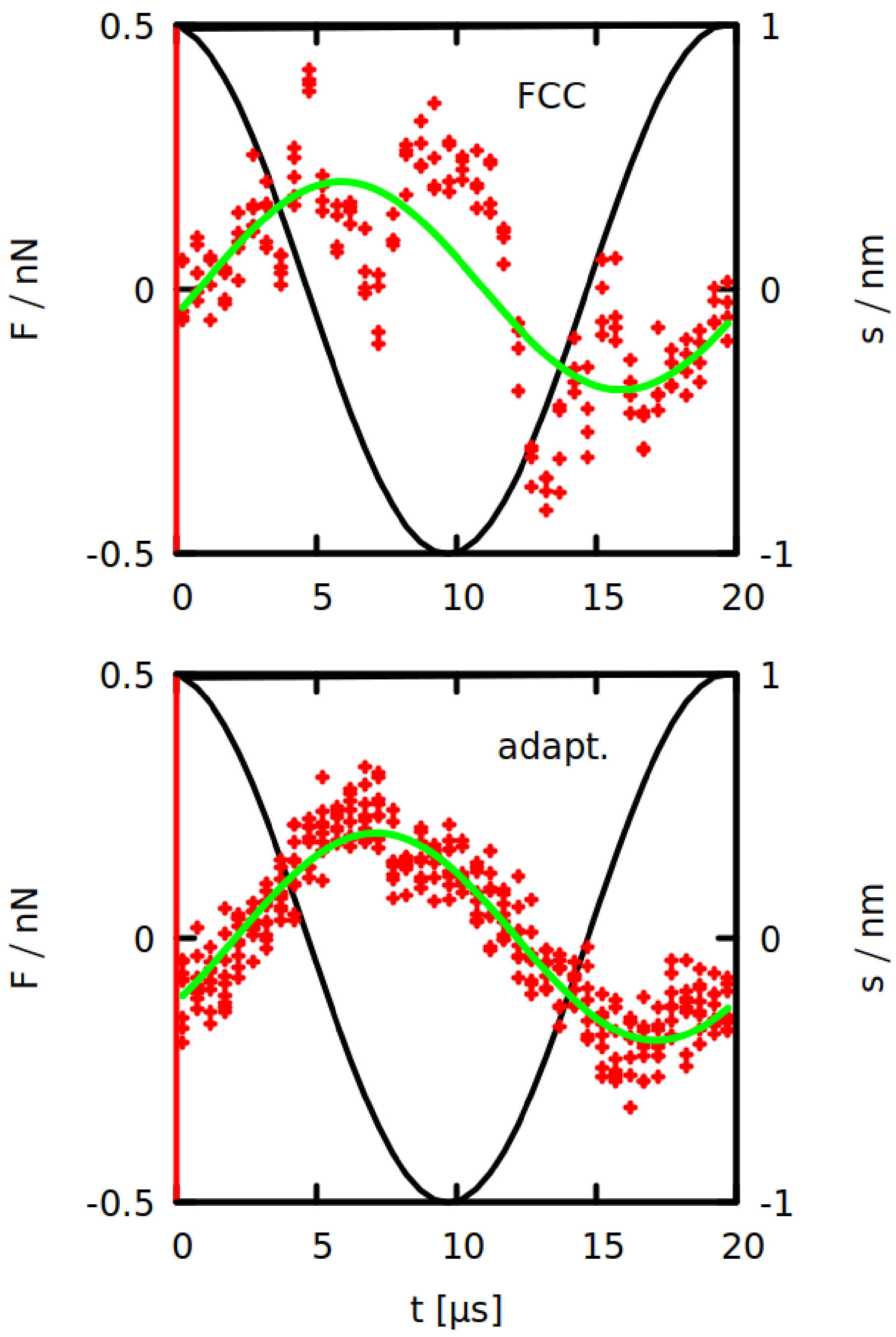


Figure S10: Force profiles for shearing simulations of gluten with period 20 $\mu$s with FCC walls (top panel) or adaptive walls (bottom panel). Each red point corresponds to one averaged value of the force. The time is shown modulo the oscillation period, so all oscillations are overlayed. Green lines show the function fitted in order to determine the shear modulus.
